# The influence of the head model on magnetoencephalography-derived functional connectivity fingerprinting

**DOI:** 10.1101/2024.08.05.606621

**Authors:** Matthias Schelfhout, Thomas Hinault, Sara Lago, Giorgio Arcara, Enrico Amico, Matteo Fraschini, Daniele Marinazzo

## Abstract

Functional connectivity (FC)-based neural fingerprinting is an approach that promises to identify and/or differentiate subjects within a cohort on the basis of the patterns of statistical dependencies between time series recorded mostly if not always noninvasively, with electroencephalography (EEG), magnetoencephalography (MEG), or functional magnetic resonance imaging (fMRI). The message is that brain activity is what differentiates subjects, or what makes a neural fingerprint “unique”. In EEG- and MEG-derived FC fingerprinting, the activity recorded at the sensors is usually projected back into cortical sources by means of an inverse model depending on head and brain shapes, sensor locations, and tissue conductivity, and further reduced in dimension to obtain time series of regional activity, used to compute FC. We think that the role of the head model in fingerprinting has been so far dismissed by means of suboptimal or incomplete tests. Here we employed a set of experiments aimed to decouple recorded activity and head model for each subject, and we found that the head model has a strong influence on both identification and differentiation.

## Introduction

Biometrics is the science of identifying people using physiological features (Ratha et al., 2001). Automated person recognition is an important technology that allows recognition of individuals through unique behavioral or physical characteristics. The most common techniques use fingerprints, face, iris or signature (Jain et al., 2008).

In recent years, there has been a great interest in the so-called brain fingerprinting. Rather than on physical/morphological characteristics of the brain, the neuroimaging community has focused on the recorded activity, also extending the focus to differentiability (how difficult it is to identify a specific individual), not without some confusion. A few years ago it was proposed that there are significant interindividual differences in patterns of statistical dependencies among neuroimaging time series (functional connectivity, FC), and that these patterns can act as a “fingerprint” that can accurately identify subjects from a large group in a repeated fMRI scanning scenario (Finn et al., 2015). Similar efforts have been made in EEG (Fraschini et al., 2015; La Rocca et al., 2014) and MEG (Colenbier et al., 2023; da Silva Castanheira et al., 2021; Sareen et al., 2021).

It’s on these latter that we will focus here, starting from the fact that the effect of anatomy has not been so far addressed with the correct tests. In (da Silva Castanheira et al., 2021) empty room data projected through the individual kernel was used. This was accompanied by contradictory statements such as: “We, therefore, tested whether such individual information unrelated to brain activity contributed substantially to individual differentiation from MEG source maps. We found that differentiation performances were considerably reduced using empty-room data (<20% across all tested models)” from which would follow that “The participant’s anatomical and head-position information embedded in their respective MEG source imaging kernels were also not sufficient to differentiate individuals.” and ultimately than “fingerprint is robust to confounders” (including anatomy). In (da Silva Castanheira et al., 2024) a linear correlation was performed “between the matching of neurophysiological profiles of twin siblings and the similarity of their respective brain anatomical features”, excluding the head, and then performing an indirect test of correlation, further giving to it a direction, concluding (“The Matching Between the Neurophysiological Profiles of Monozygotic Twins Is Not *Driven* by Anatomy “). There can be multiple pathways through which the anatomy can influence the neurophysiological identifiability which are not a direct linear relationship between the anatomical identifiability and the neurophysiological one.

Our hypothesis is that both the inverse model (projecting sensor data to sources in the brain using a kernel function based on the head model), as well as the spread of cortical activity to the sensors will lead to an influence of the anatomy on reconstructed activity and connectivity, and hence on fingerprinting. Our hypothesis stems from the evidence that electrical field spread and source mixing are not completely eliminated by source reconstruction (Anzolin et al., 2019; Hauk et al., 2022; Hincapié et al., 2017; Mahjoory et al., 2017; Palva et al., 2018; Schoffelen & Gross, 2009).

By “head model” we refer here to the conceptual and practical implementation of the anatomy of the head, relevant both in considering how electrical activity propagates from brain sources to sensors, and in the inverse modelling, when the sensor activity is projected onto brain sources.

Together with the obvious fact that the shape of the head is undoubtedly a distinctive feature, it is thus reasonable to imagine that the head model could contribute to the identifiability and differentiability of source reconstructed FC. Here we perform a series of manipulations that in our opinion constitute appropriate counterfactuals to the lack of substantial effect of anatomy, and try to provide an account of the relative influence that 1) the leadfield and 2) inversion model have on the results.

## Methods

### Data

In this paper we used resting state data from a subset of the CAM-CAN MEG dataset (Shafto et al., 2014; Taylor et al., 2017), consisting of 46 young adults (18–35 years) and 74 older adults (60–85 years). Our institutions approve (re)analysis of shared data as long as the institution that collected the data obtained approval from their ethics board.

The retrieved data were already partly preprocessed as described in (Hinault et al., 2021) using the temporal signal-space separation approach (tSSS): 0.98 correlation, 10-s window; bad channel correction: ON; motion correction: OFF; and 50 Hz +harmonics (mains) notch. Afterwards filtering with a 0.3-100 Hz bandpass was applied to ensure proper artefact detection and to more accurately identify physiological noise; subsequently, temporal segmentation into epochs, averaging and source estimation were performed. These steps were conducted in Brainstorm (Tadel et al., 2011). In addition, physiological artefacts (e.g. blinks, saccades) were identified and removed using spatial space projection of the signal. In order to improve the accuracy of the source reconstruction, the FreeSurfer (Fischl, 2012) software was used to generate cortical surfaces and automatically segment them from the cortical structures from each subject’s T1-weighted anatomical MRI. The advanced MEG model was obtained from a symmetric boundary element method (BEM model, OpenMEEG; Gramfort et al., 2010; Kybic et al., 2005), fitted to the spatial positions of each sensor (Huang et al., 1999). A cortically constrained sLORETA procedure (Pascual-Marqui, 2002) was applied to estimate the cortical origin of the scalp MEG signals, using a surface-based (one-dimensional, orientation constrained) source model, using the option to flip the sign of the sources, when required, to match the majority of the atlas region. It is important to stress at this point that when using sLORETA there is no noise covariance matrix needed, and the identity matrix is commonly used. It is worth mentioning that, as speculated in (da Silva Castanheira et al., 2021), the choice of the source reconstruction method has little effect on the fingerprinting performance. Even in the extreme choice of no regularization at all, using the leadfield only to project sensor data to the sources, the fingerprinting measures remain virtually unaltered (see Supplementary material).

The two dimensions of the kernel matrix used to perform the inverse model are the number of MEG channels and the number of reconstructed sources, here equal to 338 and 9001 respectively. Concerning the sensor number, the MEG instrument for the CamCAN dataset (Elekta Neuromag) comprises two-thirds gradiometers and one-third magnetometers: both sensor types are taken into account in source reconstruction. In Brainstorm, the “noise covariance” matrix computed from the recordings “pre-whiten” the data by pre-scaling the channels by the observed std. deviations seen in the noise data. This brings the two types of sensors into the same basic range of units. The same whitener is applied to the head modeling, to keep units and scales consistent. Source estimation then proceeds normally with a combined array of pre-whitened sensors and pre-whitened data.

The 68 time series used to estimate FC were obtained from the first mode of the principal component analysis (PCA) decomposition of the activation time course in each region of interest (ROI) from the Desikan-Killiany atlas (Desikan et al., 2006). Temporal averages of 200 ms sliding time windows (50% overlap) were then extracted across the epochs of interest.

### Measure of statistical dependency

To assess FC in the frequency domain we used the amplitude-envelope correlation (AEC). The choice was motivated by the fact that AEC is the measure used in the seminal paper of MEG fingerprinting (da Silva Castanheira et al., 2021) and the most used one in subsequent papers. In the supplementary material we consider the version with orthogonalization (Hipp et al., 2012), designed to remove the signal components with the same phase, a phase based method, the corrected imaginary part of the phase locked value (ciPLV, (Bruña et al., 2018)), as well as Pearson correlation in the time domain between wide-band signals.

### Measures of fingerprinting

The identifiability matrix, which is at the basis of the fingerprinting procedure, is a square matrix of size determined by the number of subjects. In the case of FC-based fingerprinting, each element I_i,j_ contains the values of the correlation between the vectorized upper (or lower) half of the functional connectivity matrices of subject i at time point T1 (y axis in Figure 1) and subject j time point T2 (y axis in Figure 1). This results in an asymmetric matrix, whose diagonal contains the values pertaining to the same subject at two different times (self-similarity).

**Figure 1:**
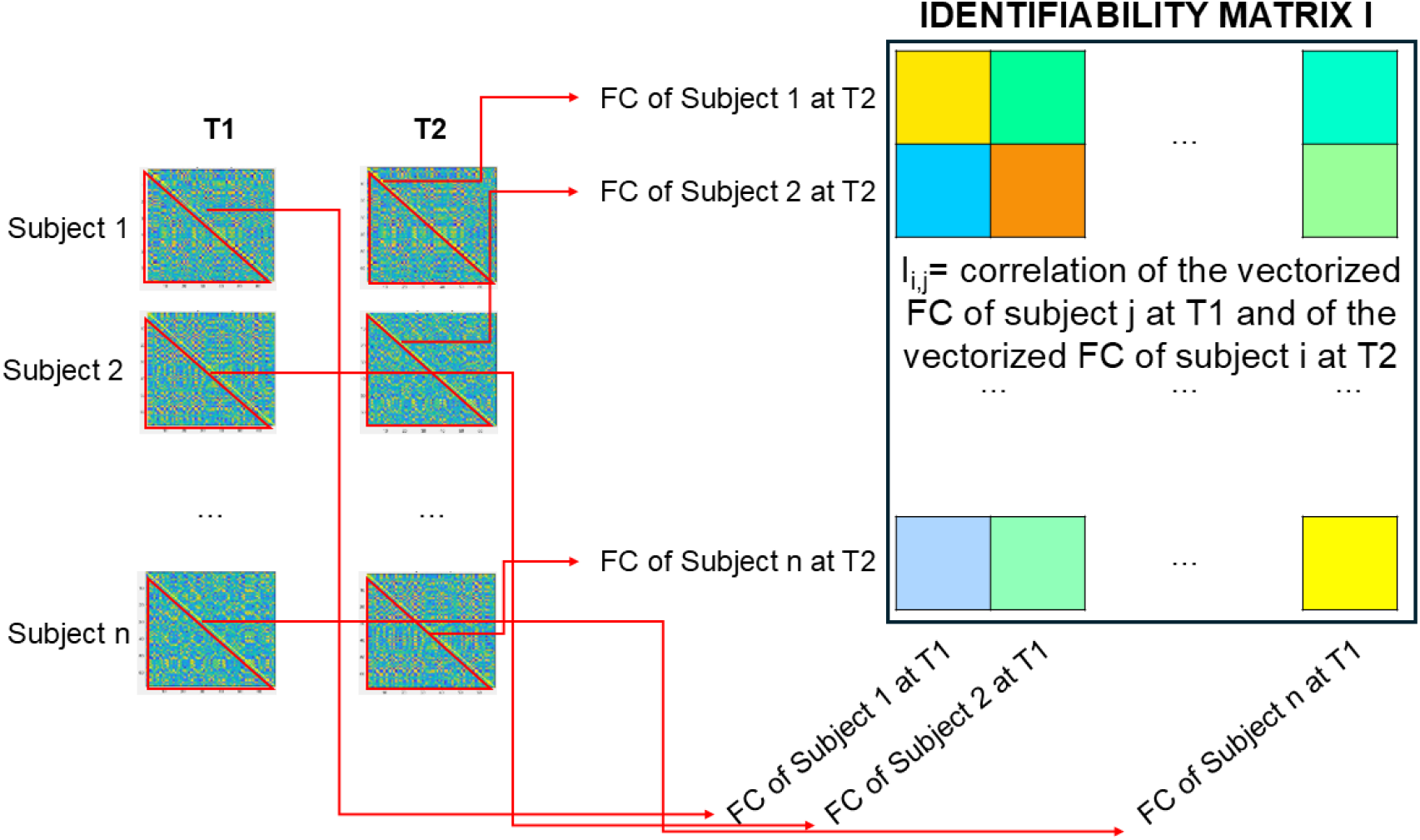
From the individual correlation matrices of n subjects at time points T1 and T2 to the identifiability matrix. There is no colorbar since this is a cartoon. When Pearson correlation is used for both FC and identifiability matrices, the values are in the range [-1 1].

From this identifiability matrix, two measures can be computed, to assess identifiability and differentiability respectively.

The Equal Error Rate (EER) is a standard identifiability measure introduced to summarise the performance of a biometric system. It relies on the basic accuracy measures for biometric verification: the False Reject Rate (FRR), that is the percentage of times that a subject is incorrectly perceived as an imposter (not themselves) by the system, and the False Accept Rate (FAR), that is the percentage of imposters which are incorrectly recognized as the original subject. The EER represents the system error when the FRR equals the FAR. Lower values of EER indicate better classification performance. These are the metrics originally used for fingerprinting studies.

Other measures were introduced in recent years, and specifically in the field of “brain fingerprinting”, with the aim of measuring “differentiability”, but sometimes still conflated with “fingerprinting” (Fraschini et al., 2025). For completeness, and to generalize our results to recent studies, we also contemplate them here.

Self-differentiability (D_self_) was defined as the correlation of one subject with themselves over different timepoints with respect to this subject’s correlation to all other subjects (da Silva Castanheira et al., 2021). In mathematical terms this becomes 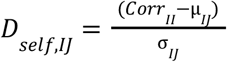, where *Corr* _*II*_ is the subject’s correlation with themselves, μ _*IJ*_ is the mean of the correlations of the subject with all other subjects, and *σ* _*IJ*_ is the empirical standard deviation. The higher the D_self_, the better this subject can be differentiated from all the other subjects. With respect to other previously introduced measures for global accuracy in differentiability such as I_self_ and I_diff_ (Amico & Goñi, 2018), D_self_ characterizes the accuracy of differentiability of one single subject in comparison to all the others.

### Our strategies for mixing/decoupling brain data and head models

We used four different strategies to test the null hypothesis of no role of the head model in fingerprinting. Below we refer to “subjects” to indicate “combination of sensor data and head model”, both in the default setting where these things belong to a single person, and in the manipulated cases where data and head models are mixed and come from different persons.

1. **Matched head model:** this is the default setting, where the head model of the same subject is used to project the sensor data to the sources, from which the functional connectivity for the subject is obtained. This is the sole case for which a “subject” in the identifiability matrix always corresponds to the same human being.
2. **Same brain data:** sensor data from a subject, not further used in the fingerprinting, are projected to the brain according to their own head model. These source reconstructed data for an external subject would represent the “brain activity” that we expect to be transferred to the sensors when recording. The projection to the sensors is done according to the head model of each of the subjects. Starting from this point, the procedure is the same as described in point 1. If “brain activity” was the sole responsible of brain FC fingerprinting, this strategy should lead to complete lack of differentiation.The first subject in the pool was chosen as the provider of the “brain activity”. Given the null hypothesis for this setting, any subject (but also random noise for what is worth) would serve the purpose.
3. **Same head model:** sensor data for each subject are projected to the source level via a single head model (of a subject not used in the fingerprinting procedure). From these sources, FCs are derived. If the head model had no effect at all, this strategy would give the same results as the matched head model. Similarly to the previous case, given the null hypothesis for this setting, any subject’s head would serve the purpose.
4. **Sensor data FC:** The source reconstruction step is omitted, and FC for each subject is obtained from the sensor data. The FC matrices have in this case a different dimension, being determined by the number of sensors rather than by the number of parcels. The identifiability matrix has on the other hand the same dimensions as in the other cases.

## Results

### EER

Figure 3 displays the EER for AEC-based fingerprinting, in five different frequency bands, for each of the four strategies depicted in Figure 2.

**Figure 2.**
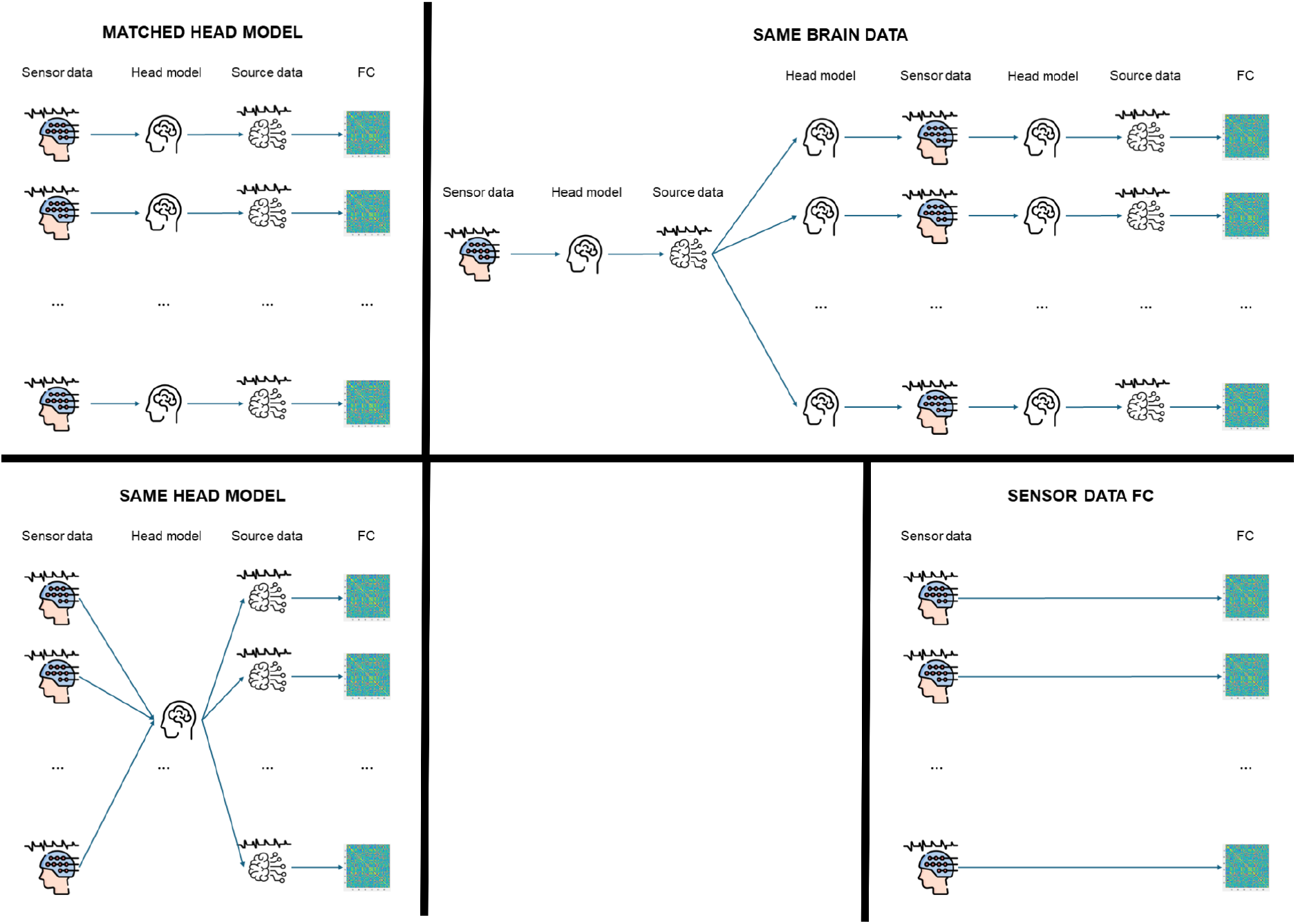
The different combinations of sensor data, head model, and source data used to obtain the Functional Connectivity (FC) matrices to be used in the fingerprinting.

**Figure 3.**
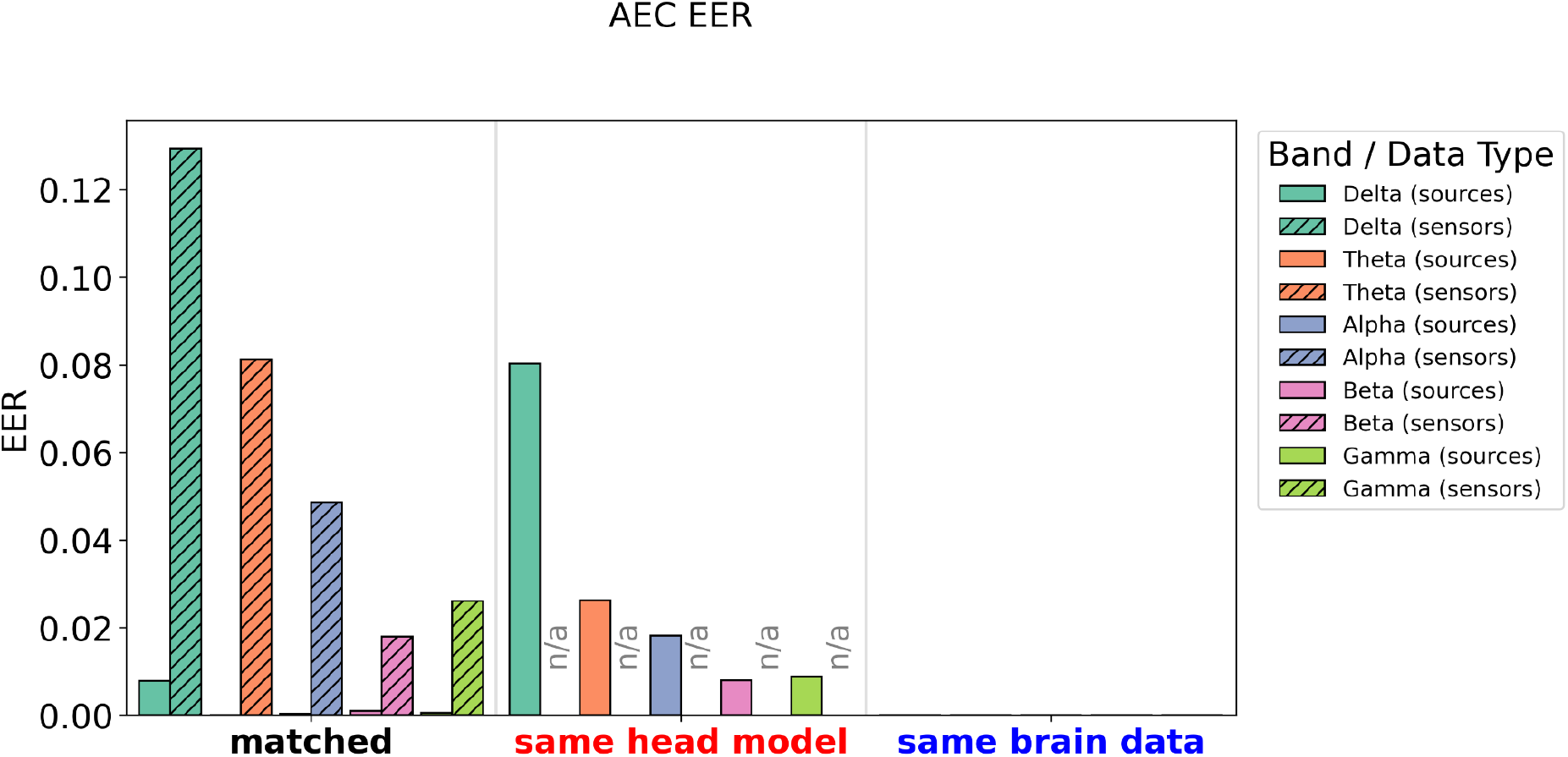
Equal Error Rate (EER) obtained from the fingerprinting based on the Amplitude Envelope Correlation (AEC), for different frequency bands, different scenarios. “n/a” indicates that the measure is not defined for that combination of data type and scenario, as opposed to instances in which the EER is zero and no bar is visible.

The first result is that sensor data worsen the identifiability, as expected and reported in previous studies.

The effects of the different scenarios are uniform across bands, with the Equal Error Rate decreasing overall with increasing frequency, and slightly increasing again in the gamma band; we will then just proceed to the description of any of the bands.

The Matched case results in excellent to perfect recognition. The performances drop in the Same Head Model case. The main results of this study are those for the Same Brain Data scenario. Here the “neural data” is the same. Yet, the identification is always perfect, thus better than in the matched case.

The supplementary material contains the results for Idiff, confirming the results above, net of the difference between what the two metrics are supposed to indicate.

### D_self_

For this measure, which is defined for each subject to indicate its individual differentiability, we report in Figure 4 the scatter plots of the value obtained with the different manipulated scenarios, indicated with “Tweaked”, against the default “Matched” setting, for the theta band. The results for other bands, and for the orthogonalized version, are reported in the Supplementary Material. The results show that for AEC the Same Head manipulation significantly reduces differentiability, similarly to what observed when fingerprinting is measured by EER. The D_self_ for the Same Brain scenario is only marginally affected.

**Figure 4.**
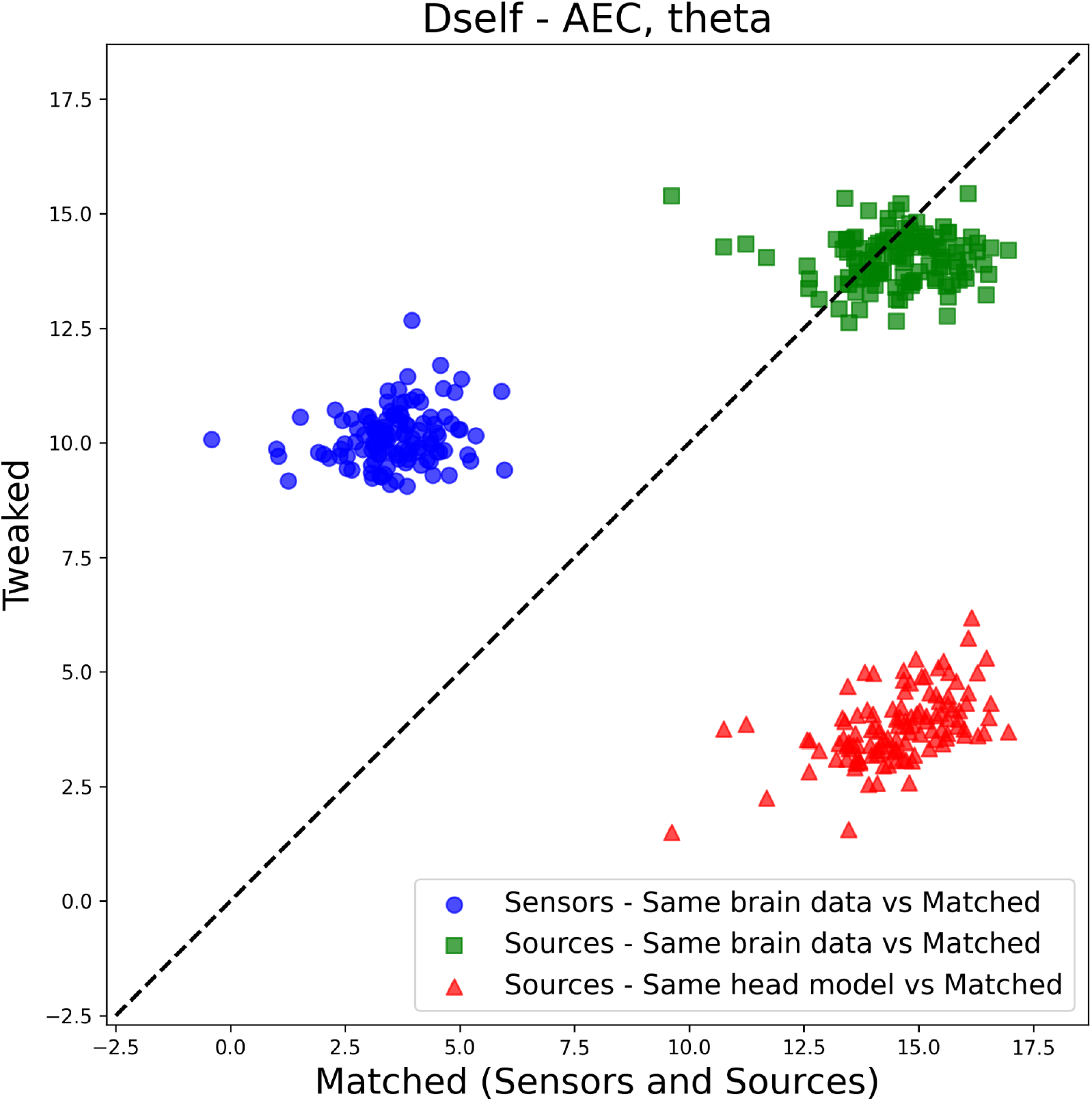
Scatter plot for AEC-based fingerprinting, of the values of D_self_ for the different manipulations of the inverse model (Tweaked) against the Matched case, for sensors and sources, in theta band. Each point represents a subject.

The attentive reader might wonder why the self differentiability improves for the same brain data with respect to the matched case when sensor data are used, while this is not the case when source reconstructed data are used. The reason rests on the fact that the departure point (the matched case) is a “bad scenario” for fingerprinting, as evident from the left panel of Figure 3, and in the dependency of D_self_ on the variance of the data. This points to a striking difference between EER and D_self_, when a low value of the former can correspond to a wide range of values of (mean values of) the latter, depending on the values of the identifiability matrix.

The averaged values for D_self_, to be compared with EER as in Figure 3, are reported in the Supplementary Material.

### Brainstorm tutorial

For improved clarity and reproducibility we modified the brain fingerprinting tutorial, which presents a reduced version of the results in (da Silva Castanheira et al., 2021) using five subjects from the OMEGA dataset available in Brainstorm (https://neuroimage.usc.edu/brainstorm/Tutorials/BrainFingerprint), to replicate the Same Head and Same Brain cases, reporting the results in Figure 5, for theta band. The results clearly reproduce the results presented in the previous sections. The subject similarity matrix (“similarity” is the name used in the Brainstorm tutorial to indicate the “D_self_” measure introduced above) for Same Head shows higher similarity off-diagonal with respect to the (Matched case. For the Same Brain case the cross-subject similarity is overall lowest.

**Figure 5.**
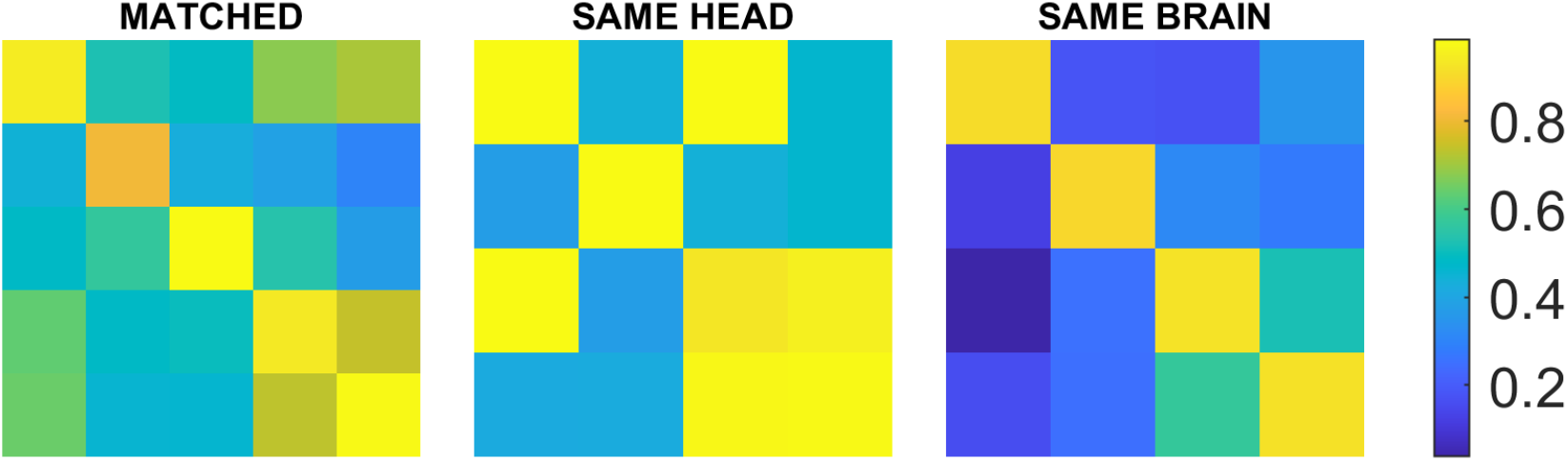
Subject similarity matrices (identifiability matrices) from the Brainstorm Brain Fingerprinting tutorial, in the original (Matched) case and in the modified ones.

## Discussion

The scope of this study was to provide a complete set of tests to establish a proper null hypothesis on the effect of the head model in the identification and differentiation of subjects (a challenge usually referred to with the broad name of “fingerprinting”) from MEG recordings. We thus applied common “fingerprinting” measures to FC matrices obtained with common measures of statistical dependency to MEG data processed according to common pipelines, combining data and kernels in different ways.

The most relevant results described in the previous sections show perfect identifiability and high differentiatiability for the Same Brain Data case, and significantly reduced performance in identifiability and differentiability for the Same Head Model case. This points to an important difference between the EER, focusing on the lack of overlap between the distributions, and the D_self_, focusing on the distance between the center of mass of the distributions, see (Fraschini et al., 2025) for detailed simulations. This behavior resonates with what reported in (Sareen et al., 2021), where the narrower distributions obtained for non-corrected measures, explained there with an increased smoothness of the activity due to leakage, allow for high separability, also in presence of reduced distance between the distributions. Likewise, the higher identifiability (lower EER) of the Same Brain Data case with respect to the matched one could be due to the fact that reduced brain data variability results in lower False Acceptance Rate, while more consistent matching scores, less likely to fall below the threshold, result in lower False Rejection Rate. These two phenomena would in turn result in reduced overlap between the distributions, and consequent lower EER. Another contributing factor could be that the “ground truth” brain activity, consisting of reconstructed source activity from a subject chosen at random, is particularly stable between the two sessions, yet the upper limit of the possible values in this case would be those of the matched case.

Amplitude-based measures like AEC, even with orthogonalization, are more sensitive to spatial leakage and head model changes because they involve signal amplitude, which does not have the phase-leakage correction inherent to phase measures such as ciPLV (Bruña et al., 2018). When the head model is shuffled or swapped, the geometry of the head and its influence on signal mixing changes. Measures that depend strongly on the reconstructed source amplitude and orientation, such as AEC, show significant performance degradation because the spatial filters applied no longer align well with the actual source orientations; Phase relationships such as the ciPLV on the other hand are overall less dependent on the head model, meaning that they are less sensitive to changes in scenarios. This can also explain why phase based measures are seldom used in electrophysiological connectome fingerprinting: their reduced dependency on the head model makes them a less convenient choice when the goal is differentiability.

With these results, we have then shown that the head model (likely through the leakage associated with both the forward and the inverse problem) is a relevant factor that influences identifiability and differentiability of a single subject. A previous seminal study on MEG-based brain fingerprinting (da Silva Castanheira et al., 2021) had tested the effects of anatomy mainly through a test consisting in projecting empty-room sensor recordings through the kernel reflecting the subject anatomy. We have two main hypotheses on the reasons behind this discrepancy. The first one is that leakage and mixing depending on the anatomy are induced in the data already at the stage of propagation of brain activity to the sensors. The second one is that the same kernels are computed using the empty room to define the noise and data covariance, and the noise covariance is used in turn to define the intrinsic noise of the sensors: this speaks to the same reasons behind the recommendations that if the target brain activity is resting, resting segments cannot be used to obtain noise statistics. On a more fundamental level, we think that starting from the same source level data (our “Same Brain” data case) represents a more appropriate counterexample for the question “are subjects differentiable by means of their brain activity?”. It could be argued that using the identity matrix for all the inversions would induce a worsening in the identification. This would mean that the “head models” (in the wide sense mentioned above) become more similar, and less characteristic. Here we would worry about the opposite effect, namely an artificial improvement of the identification. If something might worsen it, this is just a reinforcement of our results. The null hypothesis of no effect of the anatomy is rejected anyway.

Then, there’s a fundamental design reason for not using the data covariance in the kernel: when we swap head and data, we would induce a bias using the data covariance matrix (depending on the data) in the kernel, which we want to leave “head-specific”.

We then agree with (da Silva Castanheira et al., 2021) when they write “We anticipate little influence of the type of source model used though, based on evidence that beamforming kernels are mathematically equivalently to other major classes of linear source estimation kernels, such as weighted minimum-norm estimators”. We actually further pushed this claim by repeating the calculations using the leadfield only, without any regularization: the fingerprinting metrics are unchanged up to the third decimal figure (results in the Supplementary Material).

This problem is not specific to MEG/EEG. A recent study showed that brain anatomy is relevant also in shaping surface fMRI connectivity and its applications, including fingerprinting (Jeganathan et al., 2024). Some issues related to normalization to a common template were found to be of minor importance in volumetric fMRI (Finn et al., 2015).

The effect of the anatomy when fingerprinting is computed on reconstructed sources from MEG reported here should not be surprising, or not even necessarily be a bad thing. Neural activity can definitely play a role in fingerprinting, and task-induced modulations happen while the anatomy is fixed, but one cannot forget that neural activity is linked to anatomy, and some anatomical features or locations also influence the dynamics and its relevance for fingerprinting (Jeganathan et al., 2024). Not to forget that anatomy alone is itself a primary tool for fingerprinting, not only of the brain. Consequently, some interpretations in terms of “unique signature components of fast neurophysiological brain dynamics” (da Silva Castanheira et al., 2021) or along similar lines should be weighted against this evidence.

Similarly, in case of twins (da Silva Castanheira et al., 2024; Demuru et al., 2017), what is heritable and common is primarily the head shape, eventually followed by the dynamics of brain activity. The test performed in (da Silva Castanheira et al., 2024) is a linear correlation “between the matching of neurophysiological profiles of twin siblings and the similarity of their respective brain anatomical features”, first of all excluding the head, and then performing an indirect test of correlation, further giving to it a direction (“The Matching Between the Neurophysiological Profiles of Monozygotic Twins Is Not *Driven* by Anatomy “). There can be multiple pathways through which the anatomy can influence the neurophysiological identifiability which are not a direct linear relationship between the anatomical identifiability and the neurophysiological one. Here a different hypothesis is being tested, a clearer one in our opinion, with more appropriate methods (interventional/counterfactual). Again, we do not exclude all effects of something other than anatomy. The above cited paper also concludes “The neurophysiological profiles of monozygotic twin pairs match each other beyond what can be explainable by mere neuroanatomical resemblances” which is not in contrast with what is claimed here. Ultimately, similar goals could have been achieved using already established concepts such as test-retest reliability (Colclough et al., 2016). Framing these concepts as “fingerprinting” comes with some responsibilities and assumptions, which we argue that have not been properly addressed.

Another objection to the present approach could be that the goal of what has been called neurophysiological fingerprinting is not identifying subjects but rather investigating how the identifiability or differentiability are modulated by different tasks or other conditions, during which the head remains the same (da Silva Castanheira et al., 2023; Sorrentino et al., 2021; Troisi Lopez et al., 2023). We think that still in these cases the influence of the head is not irrelevant: for some subjects or brain areas more affected by the effect of the anatomy, relative change due to the task would have less weight. Here we don’t dismiss nor deny these modulations and their significance; in any case the correct statement should remain “not all effects are due to anatomy”, since there’s no direct test performed on the specific role of purely neuronal activity.

Together with the caveats exposed above, we should always be careful and align nomenclature and measurement, using consolidated performance metrics (the Equal Error Rate) when the goal is identifiability and other measures when the question is a different one (e.g. difference/separability between distributions). These two concepts are different and not univocally related, and are both affected by anatomy. Several of the metrics investigated here are not appropriate metrics for fingerprinting, as further expanded on in (Fraschini et al., 2025). We have kept those here to assess the effect of the head model in other studies where the analyses were nonetheless called “fingerprinting”.

Concerning the choice of the dataset, there was no particular reason why we chose to work with the CAM/CAN dataset for our main results, beyond the fact that it’s of good quality and publicly available, and we already had processed and analysed it in other occasions. The points that we made in this study are in our opinion general and easily exportable to other datasets. The two timepoints we considered here belong to the same session, and are spaced by 16 minutes only (as it is also the case for the Brainstorm tutorial on brain fingerprinting using the OMEGA dataset). This is something that, if anything, should favour the role of brain activity over the role of the head model, since the latter is likely more stable in time for a single subject. Finally, there are many choices, most of all valid, for data processing, parcellation, measures of statistical dependency, etc.. While any of these choices would likely be reflected in a different numerical value for the “fingerprinting” measures that we explored, we have no reason to think that the main message of this paper would change over this multiverse. The modularity of our code, and the good state of open and reproducible science in our field allows for an easy exploration of possible differences.

To conclude, the search for the stability of patterns of statistical dependencies in neuroimaging data is undoubtedly a fascinating and promising area of research. However, the current interpretation of findings is influenced by existing methodological limitations and potential confounding factors. Here we addressed a family of these, affecting MEG data. We believe that properly addressing the impact of head models and anatomical factors could serve as a crucial foundation for refining our understanding of brain fingerprinting, both in theoretical and practical contexts.

## Supporting information

Supplementary results

## Data and code availability

The raw data are available at https://cam-can.mrc-cbu.cam.ac.uk/dataset/.

Code is available at https://github.com/MatthiasSchelf/brain-fingerprinting.

Results and identifiability matrices are available at https://zenodo.org/records/12795882.

## Acknowledgements

GA and SL were supported by the Italian Ministry of Health (GR-2018-12366092 awarded to GA).

## Notes

https://cam-can.mrc-cbu.cam.ac.uk/dataset/

https://zenodo.org/records/12795882

### Competing Interest Statement

The authors have declared no competing interest.

### Summary of Updates

simplified code and results, moving things to supplementary material

