## Supplementary results for "The influence of the head model on magnetoencephalography-derived functional connectivity fingerprinting"

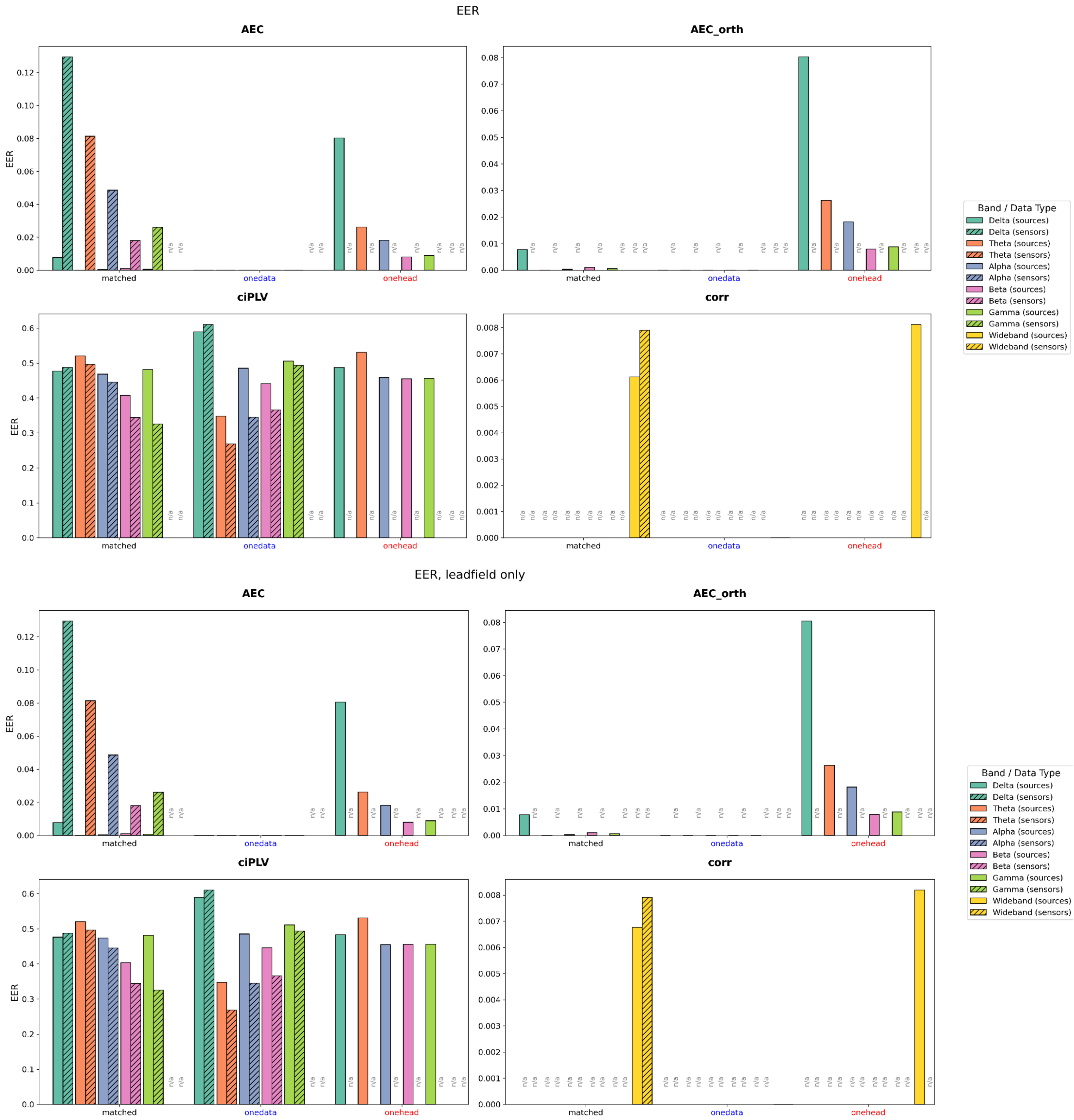


Figure S1. Equal Error Rate (EER) obtained from the fingerprinting based on four different measures: the Amplitude Envelope Correlation (AEC) with and without orthogonalization, the corrected imaginary component of the Phase Locking Value (ciPLV), and the wideband correlation, for different frequency bands, different scenarios. “n/a” indicates that the measure is not defined for that combination of data type and scenario, as opposed to instances in which the EER is zero and no bar is visible.

The upper set of four panels reports the results obtained using the identity matrix as regularization for source reconstruction. The lower set reports the results obtained using only the leadfield for source reconstruction.

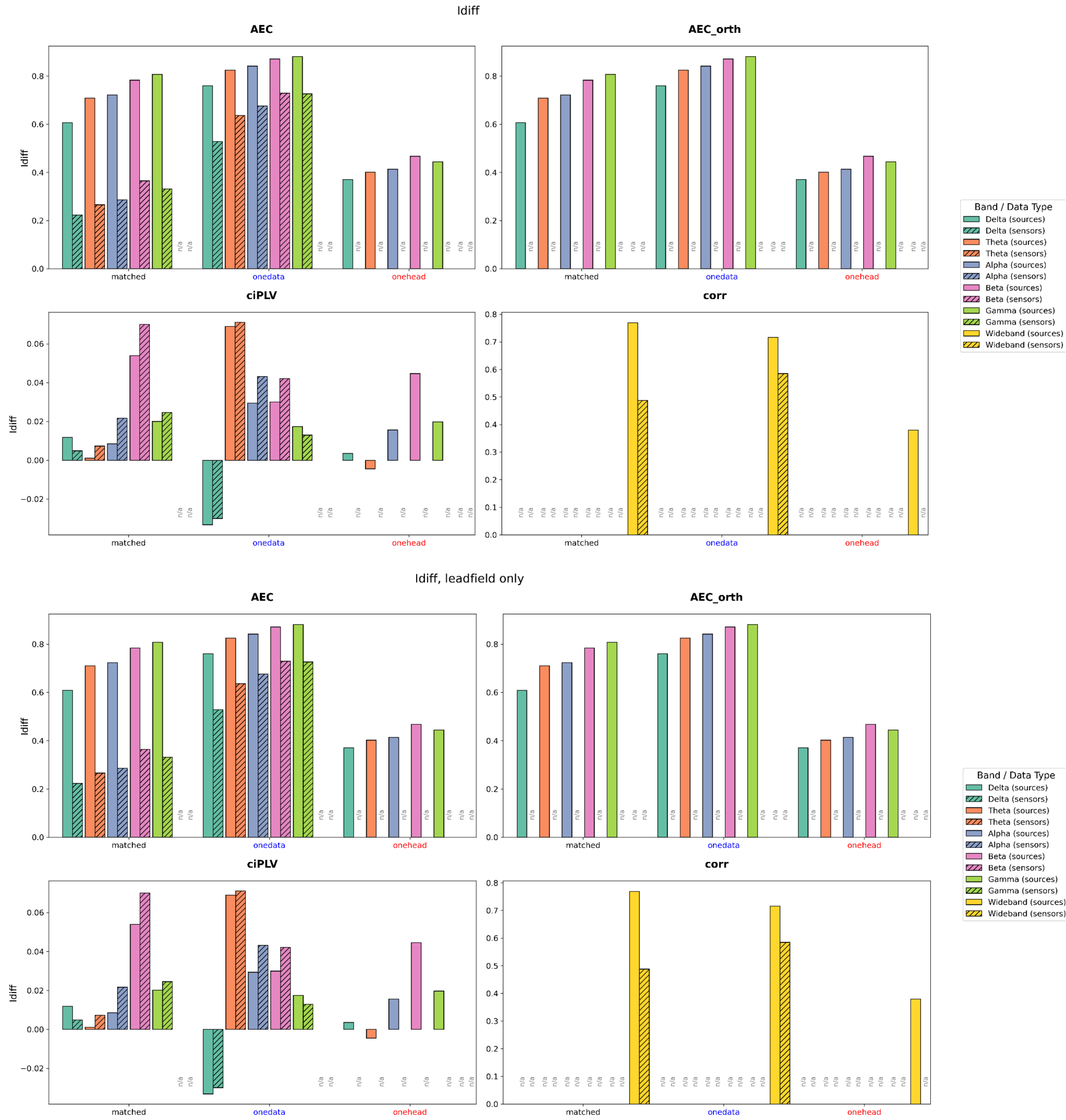


Figure S2. Differential identifiability ($I_{diff}$) obtained from the fingerprinting based on four different measures: the Amplitude Envelope Correlation (AEC) with and without orthogonalization, the corrected imaginary component of the Phase Locking Value (ciPLV), and the wideband correlation, for different frequency bands, different scenarios. “n/a” indicates that the measure is not defined for that combination of data type and scenario, as opposed to instances in which the EER is zero and no bar is visible.

The upper set of four panels reports the results obtained using the identity matrix as regularization for source reconstruction. The lower set reports the results obtained using only the leadfield for source reconstruction.

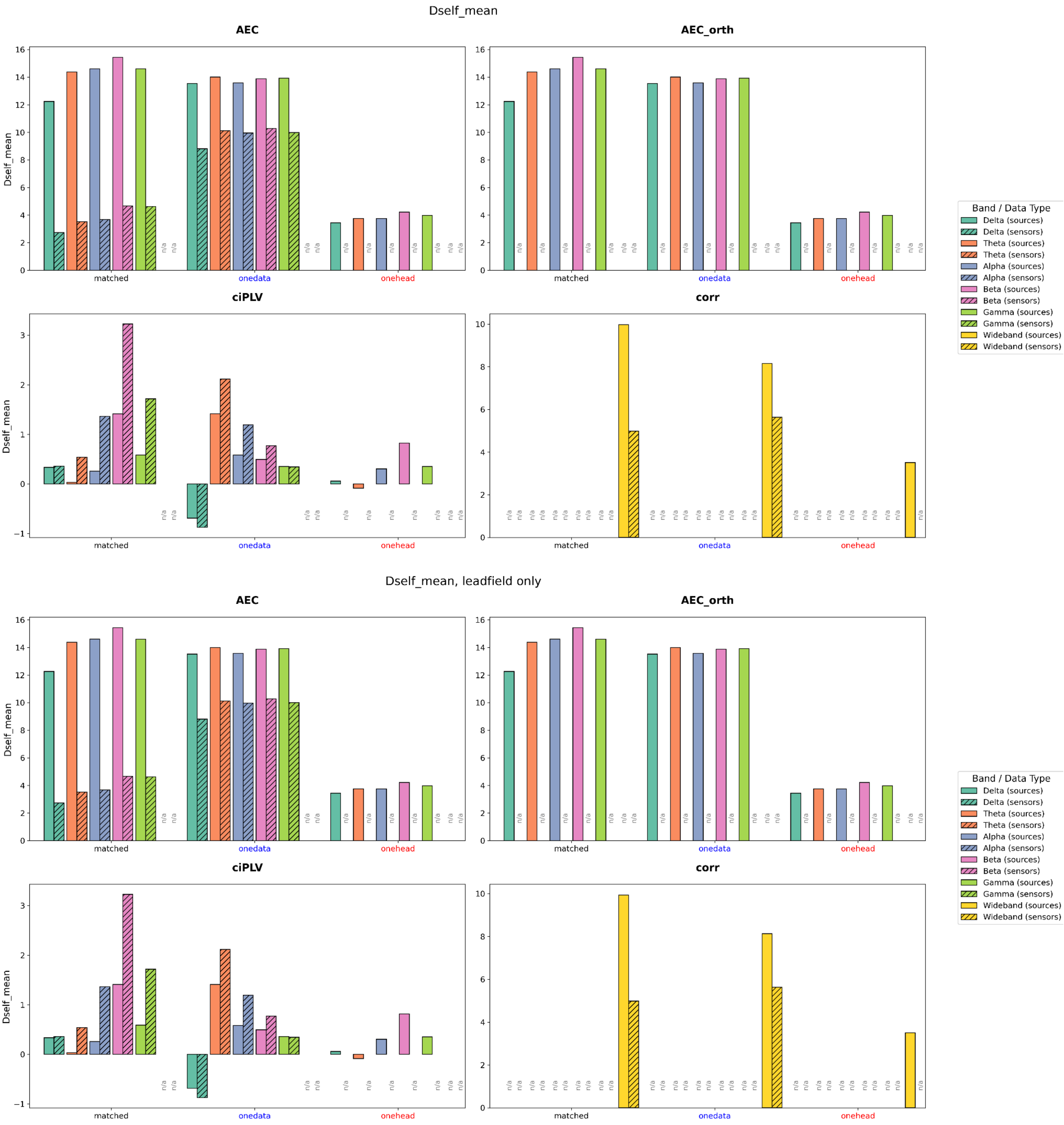


Figure S3. Differentiability ($D_{self}$) obtained from the fingerprinting based on four different measures: the Amplitude Envelope Correlation (AEC) with and without orthogonalization, the corrected imaginary component of the Phase Locking Value (ciPLV), and the wideband correlation, for different frequency bands, different scenarios. “n/a” indicates that the measure is not defined for that combination of data type and scenario, as opposed to instances in which the EER is zero and no bar is visible.

The upper set of four panels reports the results obtained using the identity matrix as regularization for source reconstruction. The lower set reports the results obtained using only the leadfield for source reconstruction.

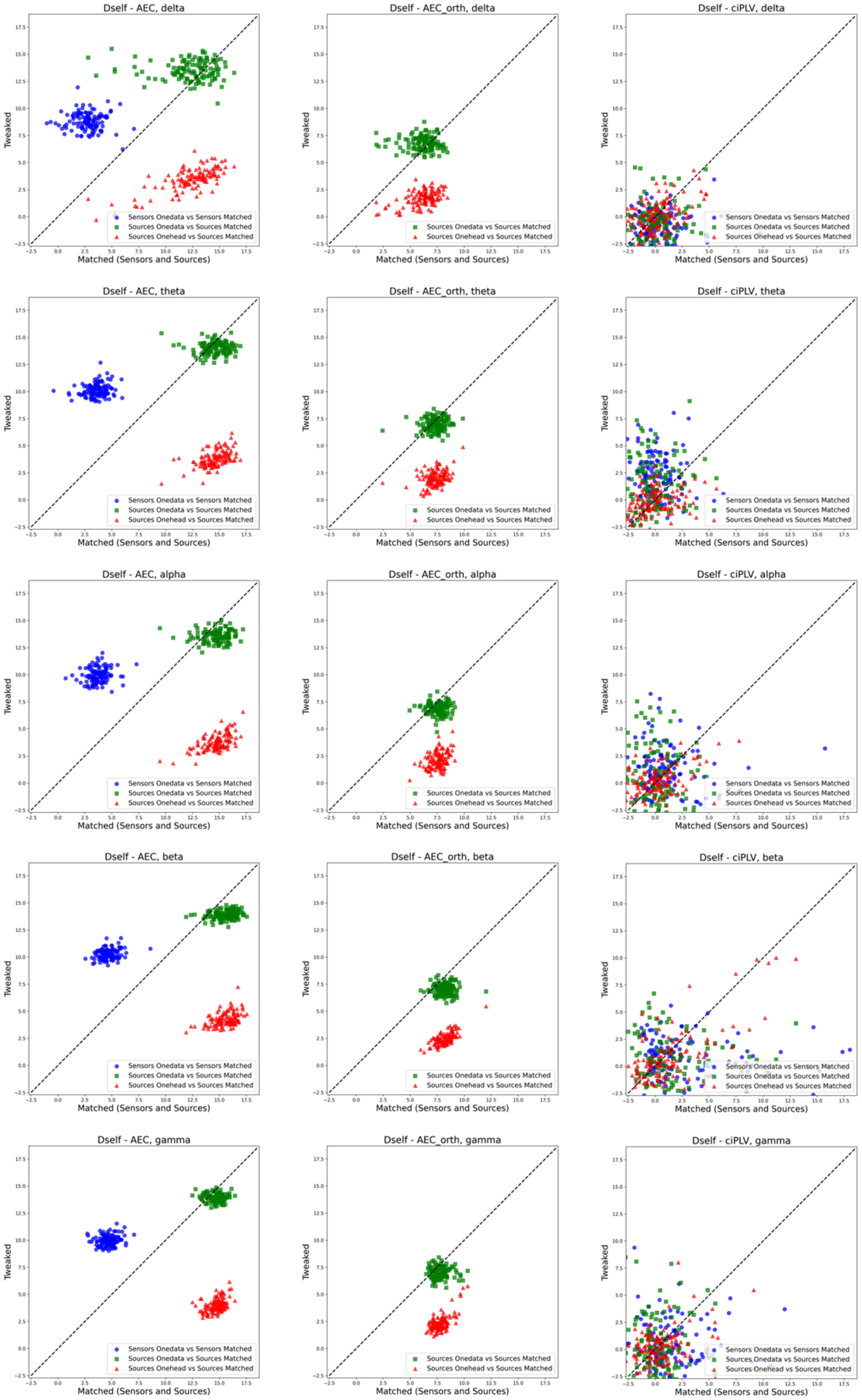


Figure S4. Scatter plots of the values of $D_{self}$ for three measures (AEC with and without orthogonalization, ciPLV) and different manipulations of the inverse model (Tweaked) against the Matched case, for sensors and sources. Each point represents a subject.
